# Expandability differences in learning transfer between computerized and non-computerized games

**DOI:** 10.1101/2020.05.14.096701

**Authors:** Maki Shigyo, Kaori Tamura, Tsuyoshi Okamoto

## Abstract

Computerized and non-computerized games are used in training designed to improve cognitive function. However, it is unclear which properties of the games influence the transfer of cognitive performance. This study aimed to examine the expandability of this transfer according to the properties of training tools. We introduced two training tools (virtual and standard Rubik’s Cubes) and examined bidirectional transfer between the two cube types and transfer to other cognitive tasks. The results showed that transfer from the virtual cube to the standard cube was greater relative to that observed from the standard cube to the virtual cube. Regarding transfer to other tasks, cognitive transfer did not differ significantly between the virtual and standard cubes, but the training exerted beneficial effects. These results suggest that transfer expandability differed between computerized and non-computerized games. The findings of the study could contribute to the provision of an effective cognitive training programme.

## Introduction

Computerized games (CGs) are popular and enjoyed as a means of entertainment by people of all ages worldwide. Numerous CGs have been developed since the 1960s, and the quality of the games’ graphics and sound has increased considerably. Originally, CGs were created to provide users with pleasure and enjoyment. However, recent reports have indicated that playing some CGs could exert beneficial effects on cognitive function, as they require concentration, memory, coordination, and rapid reaction^1^. In accordance with these beneficial effects, new CGs, commonly known as ‘brain training games’, have been designed to improve cognitive function. Note that, brain training games are not limited to CGs. Non-computerized games (NCGs: e.g., wooden blocks, puzzle rings, and clay play) have been used to improve cognitive function in young children worldwide. The beneficial effects of these games on cognitive function are well known^2–7^. The transfer of cognitive performance from game playing to general cognitive function is a critical topic in education and medicine. In these fields, easy cognitive training, which maintains motivation, is required for children, older adults, and patients with cognitive deficits. The examination of the mechanisms underlying cognitive transfer via both CGs and NCGs could contribute to the establishment of easy and enjoyable training programmes designed to improve cognitive functions.

Many studies have endeavoured to clarify the general mechanism underlying cognitive transfer via games; however, it is a complex phenomenon that is affected by numerous internal and external factors^8,9^. Transfer has not been categorized clearly, but some previous studies have proposed distinctions between different types of transfer^10^. Moreover, conventional studies have examined the transfer effects of either CGs or NCGs but have not considered both simultaneously. Beneficial transfer of cognitive skills via either CGs or NCGs has been reported in many studies involving young children and older adults^11–18^. On the other hand, the study of young and middle aged adults showed no transfer effects in other types of cognitive function following certain brain training games^19^. In the studies involving adults, the transfer effects observed in participants playing CGs or NCGs have been inconsistent. It should be noted that previous studies have focused on the content, rather than properties, of the games. Therefore, the precise properties that influence the expandability of transfer effects remain unclear. In other words, it is unknown whether CGs or NCGs produce greater expandability in learning transfer.

The objective of the study was to evaluate transfer effects according to the properties of training tools in healthy young adults. The main difference between CGs and NCGs involves spatial dimensions. In conventional CGs, objects or images are shown on the two-dimensional (2D) plane, while in NCGs, objects exist in reality, within three-dimensional (3D) space. We hypothesized that CGs would improve the extent of cognitive function and exhibit greater expandability in transfer, in accordance with the results of previous studies indicating that improvement in cognitive function occurred via certain types of training if the processes involved in both trained and untrained tasks overlapped^20,21^. Objects presented on a screen are restructured to form 3D objects in the brain when individuals play CGs in which objects are displayed on a 2D plane. Mental imagery and manipulation of 3D objects are complex, high-order cognitive processes that require cognitive ability^22^. In contrast, NCGs do not require this ability, because the 3D objects are originally recognized in 3D space. We hypothesized that there would be differences in trained cognitive function between CGs and NCGs and aimed to identify these differences.

## Results

Participants were divided into three groups; the CG, NCG, and control groups (Table 1). Training tasks using different versions of the Rubik’s Cube were assigned to the CG and NCG groups (see Methods and Figs. 1a and b). To evaluate the difference in transfer effects between the two versions of the Rubik’s Cube, we measured the periods required to solve the puzzle using the unfamiliar version of the Rubik’s Cube following training. To evaluate the transfer of effects to other cognitive tasks, we conducted three cognitive tests: the serial reaction time test, working memory test, and mental rotation test (Fig. 2).

**Table 1.**
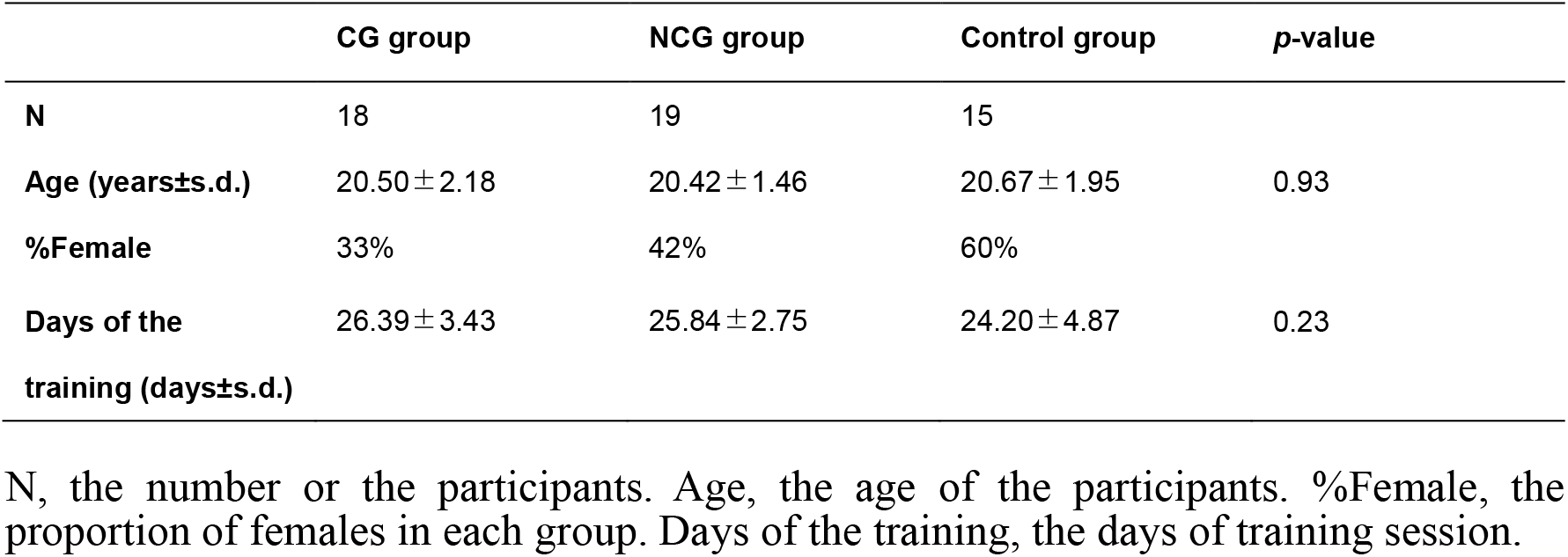
Participant information according to group.

**Fig. 1.**
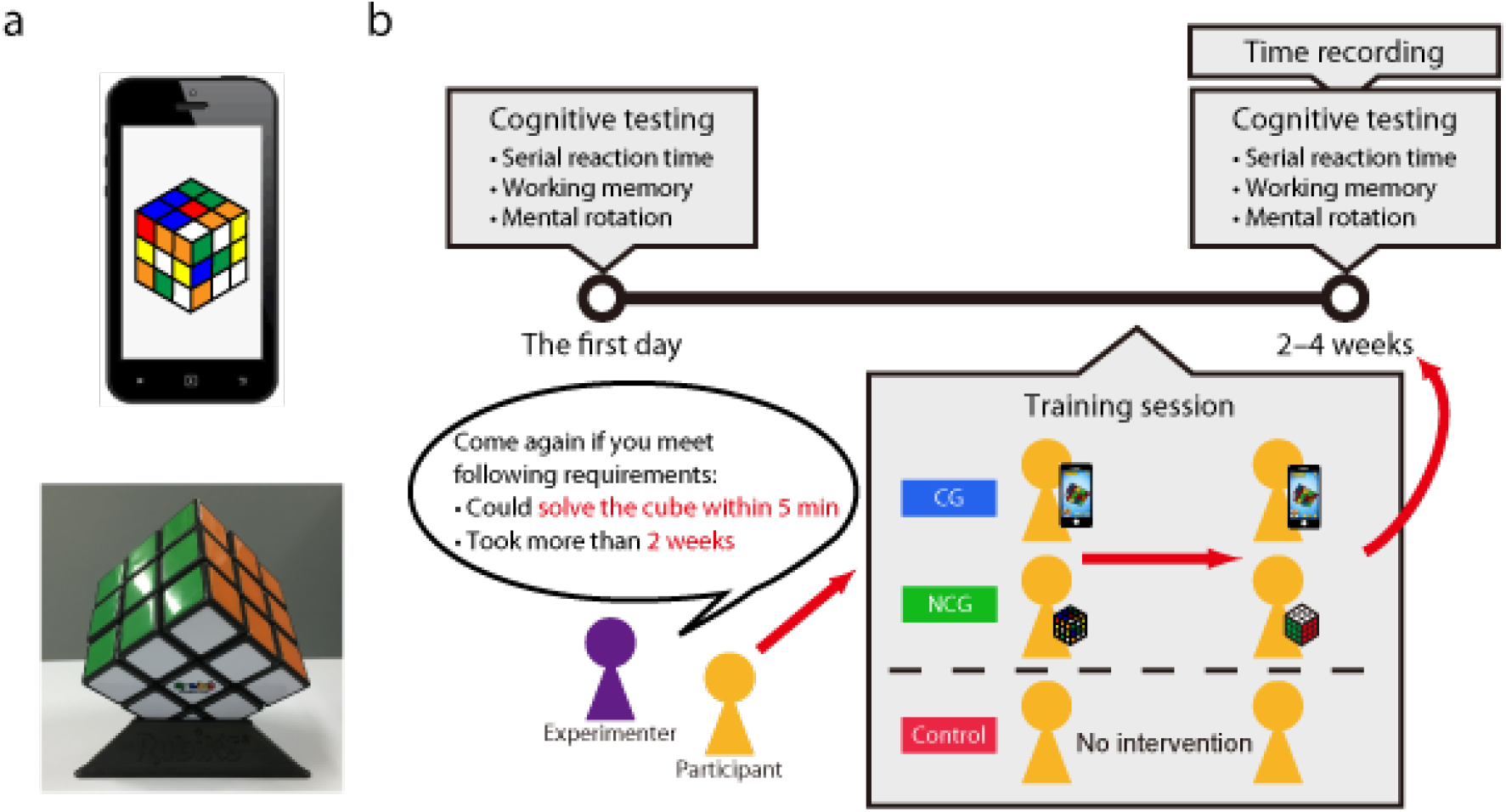
Training task and design. **a.** The study used a virtual Rubik’s Cube software application (top) and a standard Rubik’s Cube (bottom) in the training tasks. **b.**Participants in the computerized game and non-computerized game groups engaged in the training for 2–4 weeks (training session) and recorded the time required to solve the Rubik’s Cube puzzle subsequent to the training session. All participants completed cognitive tests prior and subsequent to the training.

**Fig. 2.**
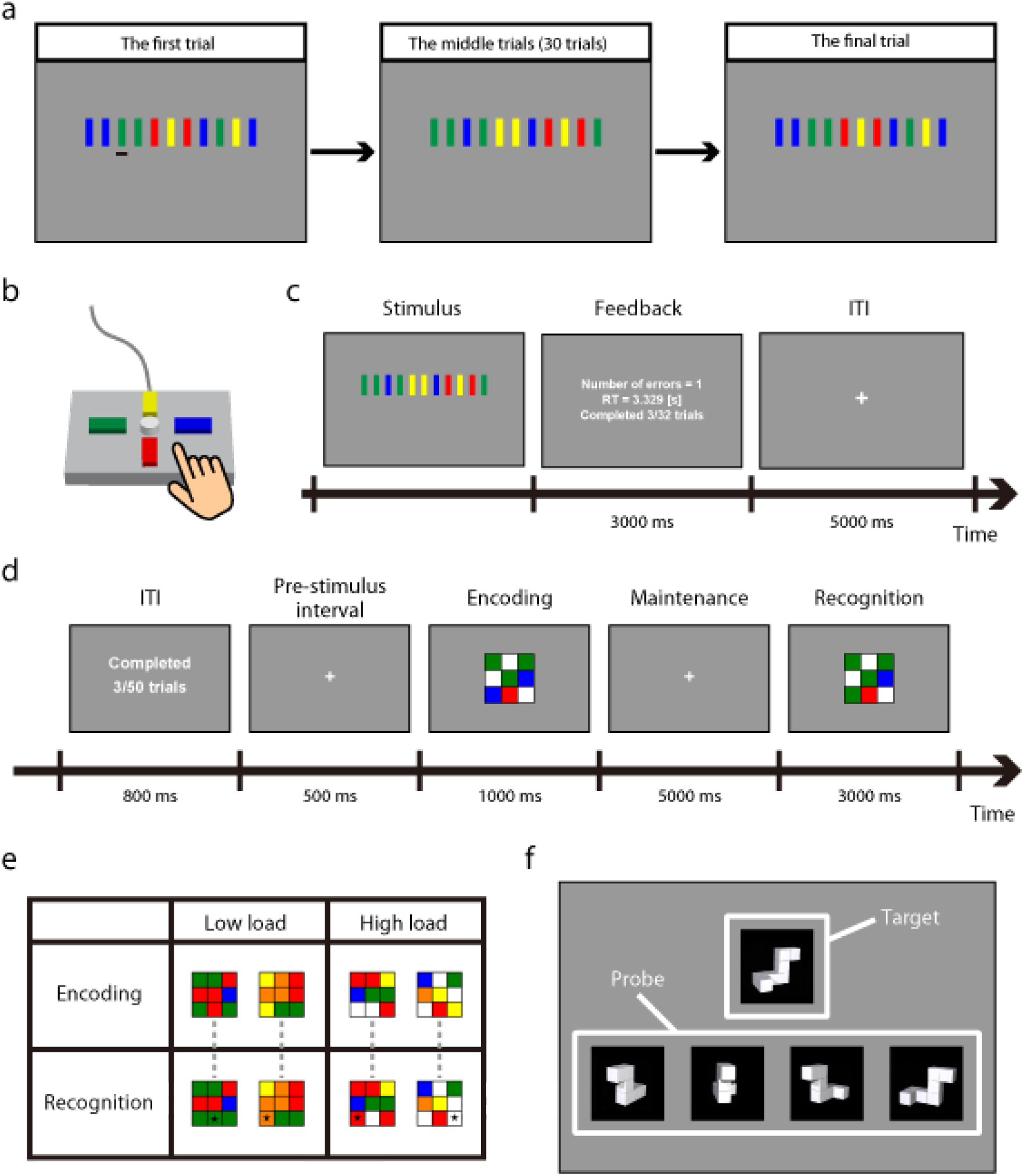
Cognitive tests. **a.** Outline of the serial reaction time test. Participants were required to use the response pad to press the button corresponding to the focal rectangle as quickly and accurately as possible. The test consisted of two stimulus sequences. One sequence was presented twice during the first and final trials, and the other was repeated 30 times successively during the middle trials. **b.** Schematic illustration of a response pad used in the serial reaction time test. The central white button on the response pad was not used in the test. **c.** The time course of a single trial in the serial reaction time test. **d.** The time course of a single trial in the working memory test. Participants were asked to identify the position of the target square out of nine small squares in the recognition stimulus, in which the colour differed from the encoding stimulus, as quickly and accurately as possible. **e.** Examples of an encoding stimulus and a recognition stimulus from both memory load levels in the working memory test. Black star marks represent correct responses. **f.** Examples of the stimuli presented in the mental rotation test. Participants were required to identify both of the standard images of the target using probe stimuli. ITI = inter-trial interval

### Bidirectional transfer between the virtual and standard Rubik’s Cubes

During training sessions, participants in the CG and NCG groups underwent training in solving the Rubik’s Cube puzzle within 5 min. The numbers of total hours required to learn how to solve the puzzle differed between participants. The average periods required to learn how to solve the puzzle within 5 min (*i.e.,* the total number of hours playing with the cube in training sessions) were 11.46 h (*SD* = 6.19) in the CG group and 15.67 h (*SD* = 10.88) in the NCG group. There was no significant difference in overall training times between the CG and NCG groups, *t*(35) = −1.50, *p* = .14 (Fig. 3a). This result suggests that the effort required to learn how to solve the puzzle did not differ between the virtual and standard Rubik’s Cubes.

**Fig. 3.**
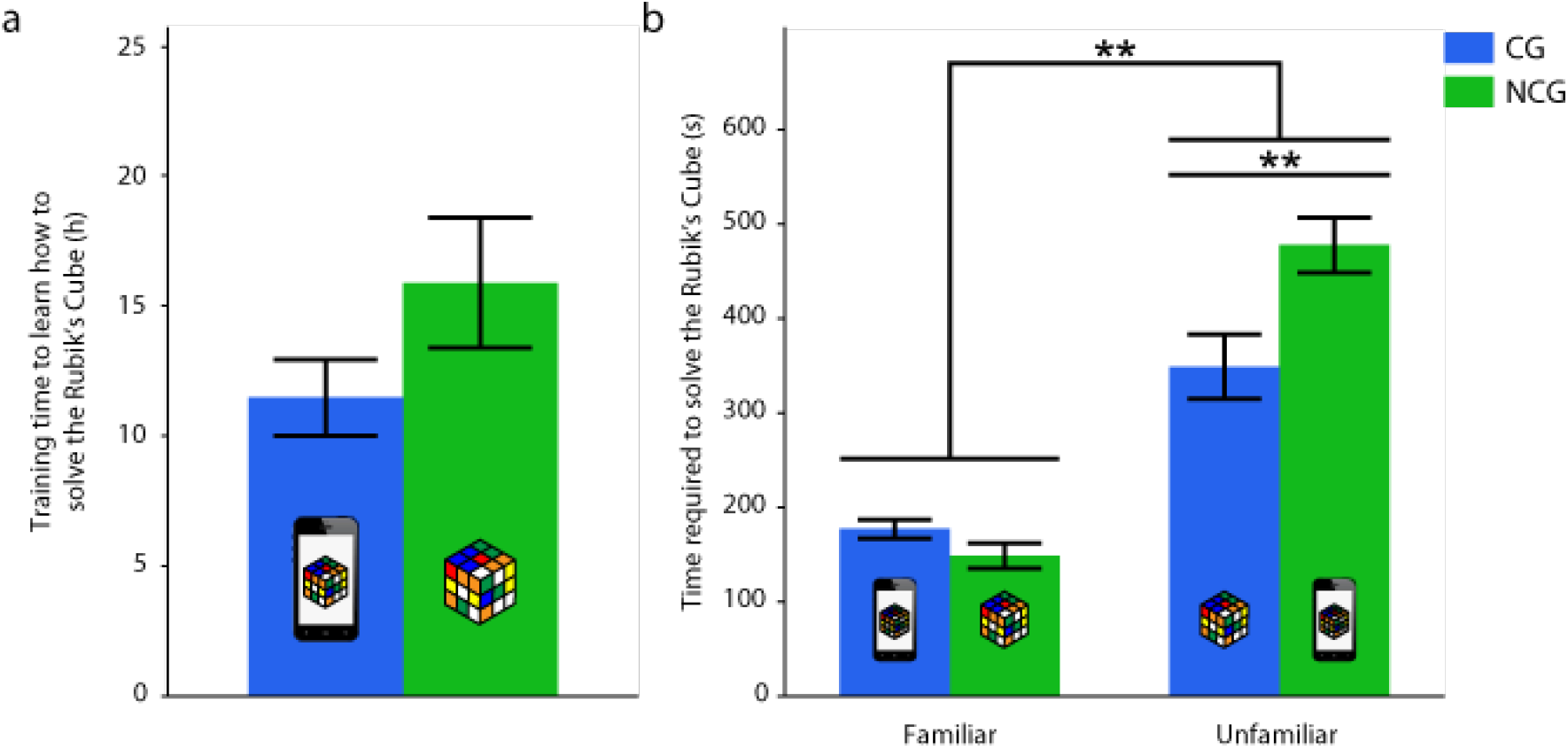
Total training time and time required to solve the Rubik’s Cube puzzle. **a.** The training time required to solve the Rubik’s Cube puzzle within 5 min. There was no significant difference in total training times between the CG and NCG groups. **b.** The time required to solve the Rubik’s Cube puzzle using the familiar or unfamiliar version of the cube. The time required to solve the Rubik’s Cube puzzle using the unfamiliar version of the cube was significantly longer relative to that required using the familiar version of the cube in both training groups. When the participants used the unfamiliar version of the cube, the time required by the CG group was significantly shorter relative to that required by the NCG group. Each icon represents the version of the cube that was used. Error bars represent the standard error of the mean. CG = computerized game; NCG = non-computerized game; \*\**p* < .01

Following the training session, we compared the periods required to solve the puzzle using the familiar and unfamiliar versions of the cube between the CG and NCG groups. The results showed that 3 of the 18 participants in the CG group and 6 of the 19 participants in the NCG group were unable to solve the puzzle using the unfamiliar version of the cube within 10 min. Both training groups required significantly longer periods to solve the puzzle using the unfamiliar version of the cube relative to those required to solve the puzzle using the familiar version of the cube, *F*(1,35) = 111.11, *p* < .01 (Fig. 3b). The main effect of Group, *F*(1,35) = 4.34, *p* < .05, and the Group × Cube interaction, *F*(1,35) = 10.92, *p* < .01, were significant. Results of post hoc *t* tests indicated that the periods required to solve the puzzle using the familiar version of the cube did not differ between the CG and NCG groups, *p* = .41, α = .025 (Fig. 3b). However, the periods required to solve the puzzle using the unfamiliar version of the cube differed significantly between the CG and NCG groups, *p* < .01, α = .025. These results suggested that the effects of training using the virtual Rubik’s Cube showed greater transfer to the standard version of the cube relative to that observed from the standard version of the Rubik’s Cube to the virtual version.

### Transfer from the virtual and standard versions of the Rubik’s Cube to other cognitive tasks

We focused on the transfer of the effects of training using the Rubik’s Cube to three other cognitive tests: the serial reaction time test, working memory test, and mental rotation test. We compared improvements in performance between the CG, NCG, and control groups in each test. Motivation scores did not differ significantly between the three groups throughout the training session, indicating that differences in performance in cognitive tests were not caused by motivation, Group: *F*(2,143) = 0.27, *p* = .77; Week: *F*(2,143) < 0.01, *p* > .99; Group × Week: *F*(4,143) = 0.27, *p* = .90 (see Supplementary Information).

The serial reaction time test was performed to evaluate participants’ ability to learn sequences in processing information and actions. The test included two types of the sequence: one was presented only twice (once during the first trial and once during the final trial), and the other was successively presented successively during the middle trials. Reaction times (RTs) were normalized according to the RTs for the first trials in the pre- and post-training tests, to compare the RT time series between the three groups. In the middle trials of the pre-training test, all three groups exhibited reductions in RT time series, and the main effect of Trial was significant, *F*(29,1392) = 33.26, *p* < .01 (Fig. 4a). The main effect of Group, *F*(2,48) = 0.32, *p* = .73, and the Group × Trial interaction, *F*(58,1392) = 0.65, *p* = .98, in the pre-training test were nonsignificant. In the post-training test, all groups exhibited reductions in RT time series, and the main effect of Trial was significant, *F*(29,1392) = 30.87, *p* < .01 (Fig. 4a). In addition, the main effect of Group was significant, *F*(2,48) = 6.45, *p* < .01. However, the Group × Trial interaction was nonsignificant, *F*(58,1392) = 0.60, *p* = .99. Post hoc *t* tests revealed that RTs in the CG and NCG groups were shorter relative to those observed in the control group throughout the middle trials (CG vs. control group: *p* < .01; NCG vs. control group: *p* < .01, α = .025). RTs in the final trial were also compared between the three groups. The main effect of Group was nonsignificant in the pre-training test, *F*(2,48) = 1.55, *p* = .22, and significant in the post-training test, *F*(2,48) = 8.02, *p* < .01 (Fig. 4b). Post hoc *t* tests showed that RTs in the training groups were shorter relative to those observed in the control group (CG vs. control group: *p* < .01; NCG vs. control group: *p* < .01, α = .025).

**Fig. 4.**
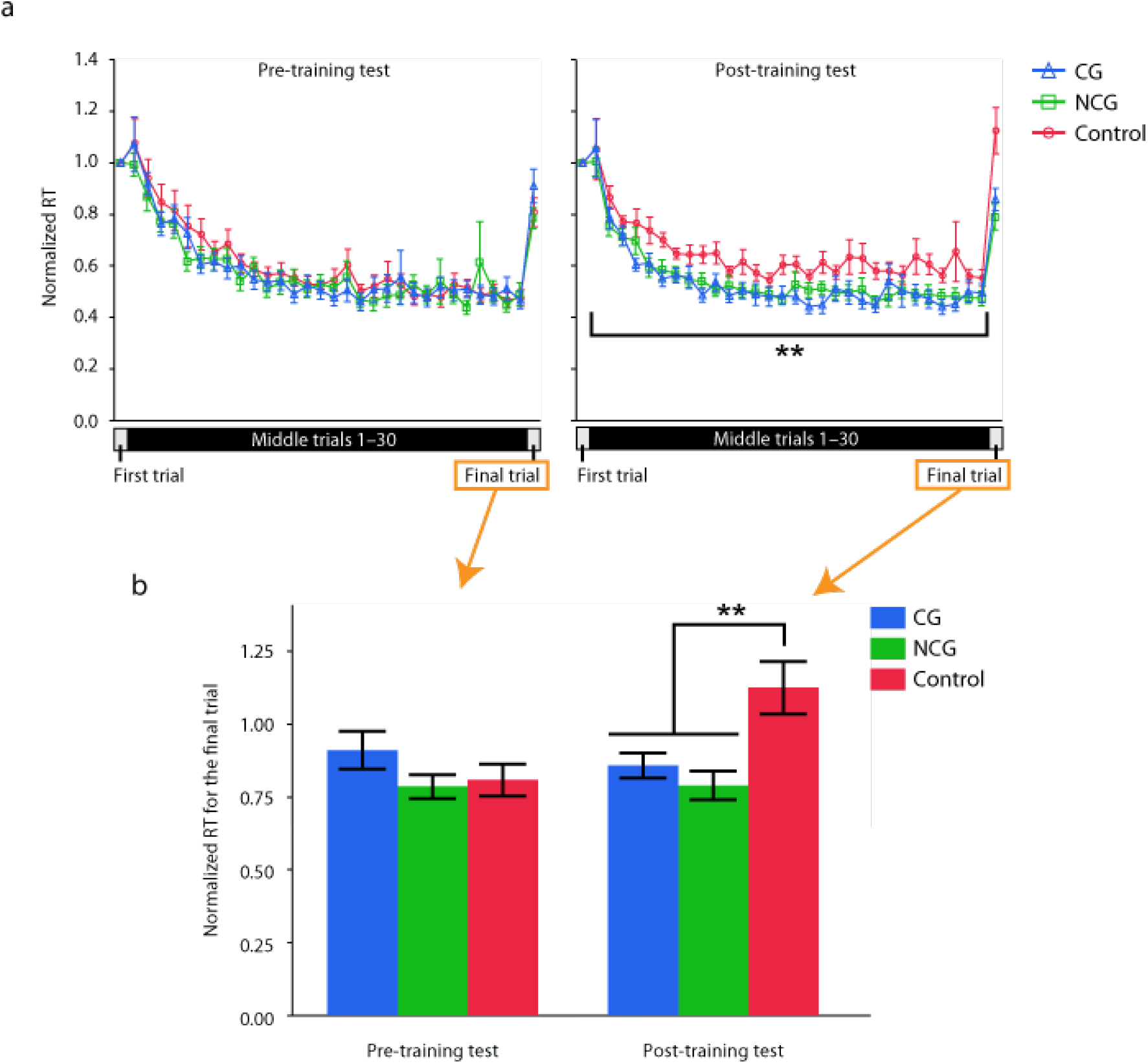
Change in performance in the serial reaction time test prior and subsequent to training. **a.** Time series for normalized RTs in the pre- (left) and post-training (right) tests. In the post-training test, the CG and NCG groups exhibited shorter RTs relative to those of the control group throughout the middle trials. **b.** Normalized RT for the final trial in the pre- and post-training tests. In the post-training test, the RTs for the CG and NCG groups were shorter relative to those of the control group. Error bars represent the standard error of the mean. CG = computerized game; NCG = non-computerized game; RT = reaction time; \*\**p* < .01.

We also analysed raw RTs prior to normalization to rule out the possibility that the results reflected a decline in the performance of the control group rather than improvements in the performance of the training groups. In the analysis of the middle trials, pre-training RTs were subtracted from post-training RTs, and the resultant RTs were compared between the three groups, using a two-way, repeated measures ANOVA. The main effect of Trial was significant, *F*(29,1392) = 3.27, *p* < .01 (Fig. 5), while the main effect of Group, *F*(2,48) = 0.53, *p* = .59, and the Group × Trial interaction, F(58,1392) = 0.61, *p* = .99, were nonsignificant. In the analysis of the first and final trials, *t* tests were performed to determine whether the RTs for the first trial differed from those for the final trial for each group and test. In the post-training test, RTs for the final trial were significantly faster, relative to those observed in the first trial, in the CG group, *t*(17) = − 3.54, *p* < .01, and NGC group, *t*(17) = −3.78, *p* < .01, but not in the control group, *t*(14) = 0.87, *p* = .40 (Fig. 5). In the pre-training test, RTs for the final trial were significantly faster, relative to those observed in the first trial, in the NCG group, *t*(17) = −5.28, *p* < .01, and control group, *t*(14) = −3.12, *p* < .01, but not in the CG group, *t*(17) = −1.62, *p* = .12. The results of the serial reaction time test suggested that training with both versions of the Rubik’s Cube increased the processing speed for sequential information.

**Fig. 5.**
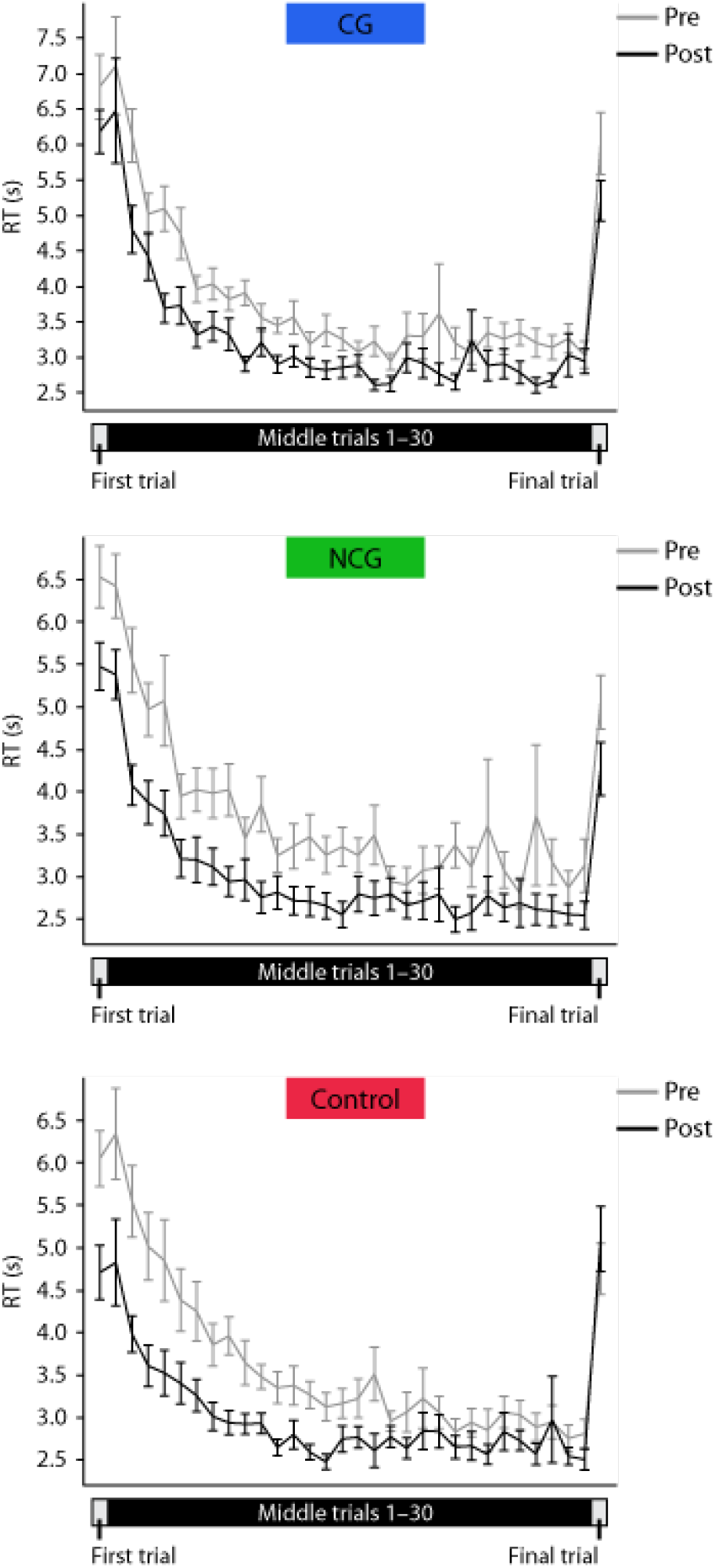
Change in raw RTs in the serial reaction time test prior and subsequent to training. Time series for raw RTs in the pre- (grey line) and post-training (black line) tests. In the middle trials, there was no significant difference in delta RTs from the pre-training test to the post-training test between the three groups. The CG and NCG groups showed significant differences in RTs between the first and final trials, but the control group did not. Error bars represent the standard error of the mean. CG = computerized game; NCG = non-computerized game; RT = reaction time

To examine the effect of Rubik’s Cube training on working memory, the participants underwent a working memory test (Figs. 2d and e). The results showed that the difference in inverse efficiency scores (IESs) between the pre- and post-training tests with high memory load were marginally significantly larger relative to those observed with low memory load, *F*(1,98) = 3.59, *p* = .06 (Fig. 6a). The main effect of Group, *F*(2,98) = 3.59, p = .22, and the Group × Load interaction, *F*(2,98) = 0.01, *p* = .99, were nonsignificant. These results indicate that improvements in performance in the working memory test did not differ significantly between the three groups.

**Fig. 6.**
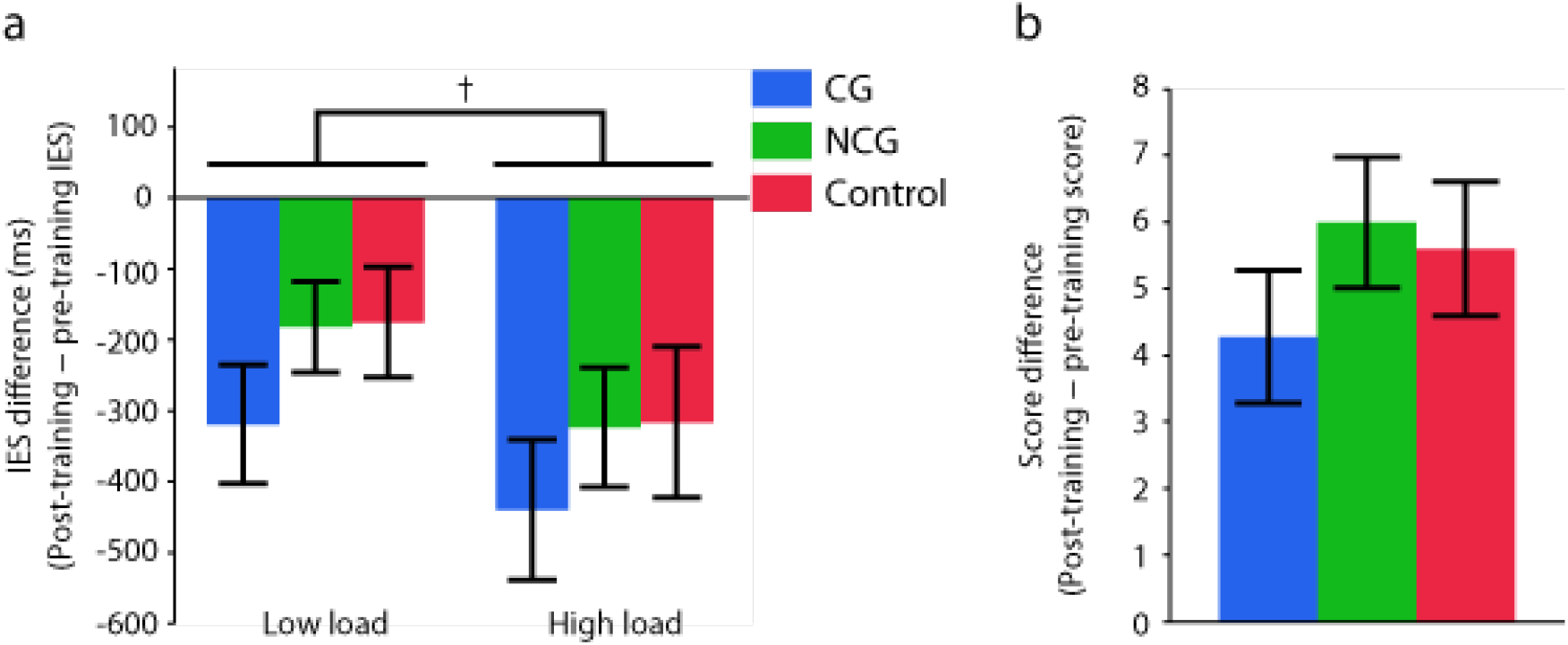
Change in performance in the working memory and mental rotation tests prior and subsequent to training. **a.** The change in IESs (post-training IES – pre-training IES) in the working memory test did not differ between the three groups for either memory load level. The change in IESs with high load was greater relative to that observed with low load, indicating a marginally significant change. **b.** The change in scores (post-training score – pre-training score) in the mental rotation test did not differ between the three groups. Error bars represent the standard error of the mean. IES = inverse efficiency score; †*p* < .10

The mental rotation test was used to assess visual spatial ability. The results showed that improvements in test performance did not differ significantly between the three groups, *F*(2,49) = 0.85, *p* = .43 (Fig. 6b).

## Discussion

The present study examined differences in transfer effects following training using a CG (*i.e.,* virtual Rubik’s Cube) or an NCG (*i.e.,* standard Rubik’s Cube). Subsequent to training for 2–4 weeks using either the standard or virtual Rubik’s Cube, participants were able to solve the puzzle using the version of the cube with which they were familiar. The time participants required to solve the puzzle using the versions of the cube with which they were familiar and unfamiliar was measured. The periods required to solve the puzzle did not differ significantly between the two training groups when participants used the version of the cube with which they were familiar. In contrast, when participants solved the puzzle using the version of the cube with which they were unfamiliar, the period required by the CG group to solve puzzle was shorter relative to that required by the NCG group. We also examined differences in the transfer of the effects of the training on other cognitive tests between the CG, NCG, and control groups. Regardless of the version of the Rubik’s Cube used, the CG and NCG groups’ post-training levels of performance in the serial reaction time test were significantly higher relative to those observed for the control group. In addition, we analysed correlations between the results of cognitive tests and experience of playing video games. The findings suggested that participants with greater experience of playing video games exhibited superior cognitive performance relative to that observed in those without this experience. In summary, the results showed that the CG provided greater expandable transfer of cognitive performance relative to that provided by the NCG.

With respect to the bidirectional transfer of effects between the CG and NCG, the transfer of the effects of using the virtual Rubik’s Cube to use of the standard version of the Rubik’s cube was greater relative to that of the effects of using the standard Rubik’s cube to the use of the virtual Rubik’s Cube, as expected. This difference in transfer effects could be explained by the difference in spatial dimensions between the virtual and standard Rubik’s Cubes. We anticipated that the participants in the CG group would be required to use mental imagery and manipulation skills to solve the puzzle. It is likely that the participants in the CG group restructured the 3D cube in their brains through the use of these skills, which were obtained via their training with the virtual Rubik’s Cube, and used this reconstruction to solve the 3D cube in 2D virtual space. In contrast, during the training with the standard Rubik’s Cube, mental imaginary and manipulation skills were not required, as participants were trained to solve the 3D puzzle in 3D real space. Nevertheless, we cannot entirely discount the possibility that the expandability of transfer effects depended on the design of the Rubik’s Cube itself. If we had used virtual reality games as training tools, the difference between CGs and NCGs would not have been observed. However, it is likely that the use of training tools in different dimensions cause differences in the expandability of transfer effects, even if the tasks involved in the training are essentially the same.

With respect to the transfer of training effects to other tasks, there was no difference in the transfer of effects to other cognitive tests between the CG and NCG groups. Moreover, we expected considerable overlap in cognitive function between the Rubik’s Cube training and cognitive tests used in the study; however, the results showed that this overlap was only slight. Some previous studies reported that transfer occurred only when the learned and transferred tasks were initially performed via similar neuronal processes^23–26^. In other words, transfer did not occur if the tasks required largely different neuronal processes. The situations involving the cognitive tests and Rubik’s Cube training appeared to overlap only slightly in the current study. Therefore, these cognitive tests did not elicit clear differences in transfer effects between the CG and NCG training, even though transfer effects were detected for both versions of the Rubik’s Cube training.

Subsequent to the training, both the CG and NCG groups showed greater improvements in performance in the serial reaction time test, relative to that observed in the control group. This finding suggests that playing with the Rubik’s Cube in both the CG and NCG groups accelerated processing speed and execution of sequential information. In other words, the participants in the training groups were able to improve their hand-eye coordination and motor learning skills for sequential information^27^. All of the participants in the training groups solved the Rubik’s Cube puzzle in accordance with the solution algorithms that they had selected. These algorithms required the memorization of sequential information to solve the Rubik’s Cube puzzle (e.g., F-R-U-R’-U’-F’, whereby a letter alone represents clockwise rotation on the given side, while a counter-clockwise turn is marked with a prime). Unlike the control group, the training groups involved the need to learn cognitive skills via which to process and execute sequential information as rapidly as possible, and participants performed the task that required these cognitive skills repeatedly for longer than 2 weeks.

In contrast to the findings regarding the serial reaction time test, improvements in performance for the working memory and mental rotation tests did not differ between the training groups and control group. One possible reason for this discrepancy is that the cognitive functions required in these tests differed from those involved in the training session. With respect to the working memory test, there are two possible reasons for the lack of transfer effects: one involves a difference in the amount of time for which the maintenance of information was required, and the other involves a difference in the content that was memorized. In the working memory test, participants were required to memorize and maintain a colour arrangement, which was similar to a side of a Rubik’s Cube, for at least 5,000 ms. However, when they solved the Rubik’s cube puzzle, they memorized and maintained the relevant information for a much shorter period, because they were required to solve the puzzle as rapidly as possible.

With respect to content, participants memorized the algorithms gradually during training. In fact, they memorized the colour arrangement only a few times while solving the puzzle, after they had memorized the algorithms. Memorizing the algorithms to solve the puzzle is thought to have allowed participants to focus on twisting the cube in accordance with the algorithms, without memorizing the cube’s colour arrangement each time. In addition, participants were required to recognize the shape of 3D objects accurately in the mental rotation test.^28^ However, they would have recognized the spatial colour distribution of the Rubik’s Cube while solving the puzzle. The Rubik’s Cube is believed to enhance spatial ability; however, there are various types of spatial ability, and it is unclear which types are affected^29^. In the current study, it is likely that solving the Rubik’s Cube puzzle required spatial distribution functions rather than mental rotation skill. Use of an appropriate test to evaluate spatial colour distribution would allow the detection of improvements in spatial ability during Rubik’s Cube training. Therefore, we considered the reason for the finding that improvements were not observed in the working memory and mental rotation tests a methodological problem rather than one involving a lack of training effects.

The results of the study provided important insight into the difference in transfer effects between two different tools, namely the virtual and standard Rubik’s Cubes. However, there is room for doubt as to whether the results reflected the bidirectional transfer of learning between familiar and unfamiliar training tools, or the original levels of difficulty involved in using the tools. The difficulty involved in the use of each tool should be measured to remove this doubt, but it would be almost impossible to record the time required by a naive participant to solve the puzzle using both versions of the cube as a control time, because by definition, a naive participant would be unable to solve the puzzle without training. Therefore, instead of recording a control time with a naive participant, we demonstrated that there was no significant difference in the total training time required to solve the puzzle between the CG and NCG groups (Fig. 3a). This allowed us to assume that the difficulty involved in solving the puzzle did not differ between the two versions of the cube. Although continuous progress monitoring during the training sessions would have provided a more direct evaluation of the difficulty involved in using each version of the cube, this type of monitoring requires precise, continuous recording of the entire training procedure, which would restrict participants’ daily activities considerably and could decrease the effects of learning transfer.

In conclusion, the results showed a difference in the expandability of learning transfer between the CG and NCG. The study provided the following two novel findings: 1) The comparison of bidirectional transfer of effects showed that the extent of the transfer of the beneficial effects from the use of the virtual Rubik’s Cube to the standard version of the cube was greater relative to that from the standard Rubik’s Cube to the virtual version of the cube. 2) The effects of training using a Rubik’s Cube transferred to other cognitive tasks in healthy young adults. We expect these findings to contribute to the provision of a substantially effective brain training programme for people of all ages.

## Methods

The study was designed to compare transfer expandability between CGs and NCGs. Participants were assigned to one of the three groups pseudo-randomly: the CG group played a CG during training, the NCG group played an NCG during training, and the control group did not undergo training (Table 1). All three groups completed identical cognitive tests at two time points, and the CG and NCG groups completed additional training tasks (Fig. 1a). The CGs and NCGs used in the cognitive training involved a virtual Rubik’s Cube, which was used via a software application, and a standard Rubik’s Cube, respectively (Fig. 1b). Rubik’s Cube (MegaHouse Corporation, Tokyo, Japan) is a popular 3D puzzle that was invented by Ernő Rubik in 1974. Both the virtual and standard Rubik’s Cubes were readily available. To determine the extent of the transfer expandability for each tool, we examined the degree of transfer between the virtual and standard Rubik’s Cubes by measuring the time required to solve the puzzle using the unfamiliar version of the Rubik’s Cube following training *(i.e.,* solving the standard Rubik’s Cube puzzle following training via the virtual Rubik’s Cube vs. solving the virtual Rubik’s Cube puzzle following training via the standard Rubik’s Cube). To determine whether the extent of transfer expandability differed between the two versions of the Rubik’s Cube, we assessed participants’ cognitive skills via three cognitive tests, which were administered prior and subsequent to the training (Fig. 2). In addition, we performed supplementary analyses to examine the correlation between cognitive performance and experience of playing video games, to confirm the results of previous studies indicating that playing video games improved some types of cognitive function (Supplementary information). All methods of this research were performed in accordance with the guidelines approved by the ethics committee at the institution with which the authors were affiliated.

### Training tools

The Rubik’s Cube, which is a puzzle game, was used in the training task. The nature of the cube allows it to be twisted and turned without breaking or falling apart. Each side of the cube is divided into nine squares of equal size. In its unscrambled state, the nine squares on each side of the cube consisted of nine square stickers of one colour. The colours of the sides of the cube are as follows: white is opposite yellow, blue is opposite green, and orange is opposite red, and the red, white, and blue sides are arranged clockwise around an apex. When people begin to play with the cube, they twist and turn it randomly, which scrambles the coloured squares. The object of the puzzle is to return it to its original form by twisting and turning it repeatedly. If they accomplish this, the Rubik’s Cube puzzle is solved.

The current study included not only the standard Rubik’s Cube (MegaHouse Corporation, Tokyo, Japan) described above but also the virtual Rubik’s Cube application developed by the MegaHouse Corporation, for use in cognitive skill training. In the CG group, the Rubik’s Cube application was installed onto participants’ smartphones or tablet-type devices, and participants in the NCG group were temporarily provided with a standard Rubik’s Cube. The CG group attempted to solve the puzzle using the virtual Rubik’s Cube, and the NCG group attempted to solve the puzzle using the standard Rubik’s Cube.

### Participants

Fifty-five healthy university students (mean age = 20.53, *SD* = 1.93 years; 24 women, 31 men) were recruited to participate in the study. All of the participants had normal or corrected-to-normal vision, and six were left-handed. Three participants who were unable to perform the training session were excluded from the study. Ultimately, the study included 52 participants (mean age = 20.52, *SD* = 1.84; 23 women, 29 men), five of whom were left-handed. All participants received a thorough explanation of the experimental procedure and provided informed written consent. Prior to commencement of the study, participants answered a simple questionnaire to determine how much time they spent playing video games, working at a desk, and exercising, and. The questionnaire included the following six questions: 1) How often do you play video games? 2) How often do you work at a desk? 3) How often do you engage in exercise? 4) When did you first start playing video games habitually? 5) When did you first start working at a desk? 6) When did you first start engaging in exercise habitually? Responses to the first three questions were provided using a scale ranging from 0 (infrequently) to 5 (frequently), and responses for the remaining three questions were provided using a scale ranging from 0 (never) to 5 (more than 12 years ago or in early childhood). The scores for Questions 1) and 4) were summed to provide a measure of participants’ experience of playing video games, the scores for Questions 2) and 5) were summed to provide measure of participants’ experience of working at a desk, and the scores for Questions 3) and 6) were summed to provide measure of participants’ experience of engaging in exercise.

The 52 participants were divided into the following three groups: the CG group (n = 18; mean age = 20.50, *SD* = 2.18 years; 6 women, 12 men; 4 left-handed individuals), which was instructed to use the official virtual Rubik’s Cube application for training; the NCG group (n = 19; mean age = 20.42, *SD* = 1.46 years; 8 women, 11 men; 1 left-handed individual), which was instructed to use the standard Rubik’s Cube for training; and the control group (n = 15; mean age = 20.67, *SD* = 1.95; 9 women, 6 men; all right-handed individuals), which was instructed not to play with any version of the Rubik’s Cube. The numbers of participants in the CG and NCG groups were higher, relative to that in the control group, because the participants in the training groups (*i.e.,* the CG and NCG groups) were expected to fall behind easily during training. Age, sex, and the days on which training sessions occurred were balanced across the three groups (Table 1). With the exception of one participant in the NCG group, the participants had little or no experience of using a Rubik’s Cube prior to the study. The participants who completed the training and the two cognitive tests received a reward for their participation, based on their performance.

### Experimental procedure

All three groups completed the same cognitive test prior and subsequent to the training sessions, which were implemented for 2–4 weeks (*i.e.,* pre- and post-training tests; for details see Fig. 2 and the Cognitive tests section below). The training sessions differed between the two training groups. The CG group engaged in training using the virtual Rubik’s Cube, while the NCG group used the standard Rubik’s Cube. Participants in the training groups were provided with training manuals that contained instructions explaining how the Rubik’s Cube puzzle could be solved using the BookLooper system (KYOCERA Communication Systems Co., Ltd., Kyoto, Japan), but they were also allowed to locate alternative solutions via the Internet. Participants were advised that they could play with the Rubik’s Cube anywhere and at any time during the predetermined training sessions. Once participants had engaged in training for >2 weeks, they were instructed to repeat the cognitive tests when they were able to solve the puzzle within 5 min without cheating. On the day on which the post-training tests were completed, the periods required to solve the puzzle using the versions of the cube with which participants were familiar (*i.e.,* the CG group solved the puzzle using the virtual Rubik’s Cube, and the NCG group solved the puzzle using the standard Rubik’s Cube) and unfamiliar (*i.e.,* the CG group solved the puzzle using the standard Rubik’s Cube, and the NCG group solved the puzzle using the virtual Rubik’s Cube) were recorded. Measurement of the time required to solve the puzzle using the virtual Rubik’s Cube was performed using a 9.7-inch iPad Air (Apple Inc., Cupertino, CA, USA), and the cube was automatically scrambled via the software application. When measuring the time required to solve the puzzle using the standard cube, the experimenter scrambled the cube based on an official scrambling programme (TNoodle-WCA-0.10.0). When the participants solved the puzzle using the cube with which they were unfamiliar, a time limit of 10 min was established. If participants could not solve the puzzle using the unfamiliar cube, the time was recorded as 600 s for convenience. Participants in the control group were instructed to spend their time as usual during the training sessions and were not permitted to play with any version of the Rubik’s Cube. Once per week, all participants were asked to provide a motivation score and record the total numbers of hours they spent playing video games, working at a desk, and engaging in exercise, using an e-learning system (customized Moodle system) during the training session. In addition, the CG and NCG groups were required to record the total number of hours spent training per week.

### Cognitive tests

To assess the transfer of cognitive skills gained through solving the Rubik’s Cube puzzle to other cognitive tasks, the participants performed the following three pre- and post-training cognitive tests: serial reaction time, working memory, and mental rotation. The order of these tests was randomized between participants but remained the same in each participant’s pre- and post-training tests. The presentation of stimuli and collection of behavioural responses were controlled using PsychoPy v1.84.2, and all visual stimuli were presented against a grey background on a 32-inch, 1,920 × 1,080 pixel display++ LCD Monitor (Cambridge Research Systems Ltd, Rochester, UK). All participants were assessed individually in a dark room, sitting approximately 114 cm from the monitor. The tests lasted approximately 30 min, and participants took a short break between tests. Prior to each test, participants received an explanation regarding the test and performed practice trials (serial reaction time and working memory tests: five trials; mental rotation test: three trials) using stimuli that were not used in the experiment, and ‘correct’ or ‘incorrect’ was displayed as feedback for each response.

The serial reaction time test designed by Trapp et al.^30^ was modified for use in the study to evaluate participants’ ability to learn sequences in processing information and actions. A stimulus sequence consisted of an array of 11 rectangles filled with one of the following four colours: yellow, green, blue, and red (3.67° × 0.93° visual angle for each rectangle, spaced evenly between 0.88° and 19.47° from each other; Fig. 2a). In this test, participants’ responses were recorded using a Cedrus^®^ response pad (model RB-530, Cedrus Corporation, California, USA), with four coloured buttons corresponding to the stimuli: yellow at the top, green on the left, blue on the right, and red at the bottom (Fig. 2b). Participants were required to press the button corresponding to the colour of the focal rectangle as quickly and accurately as possible with only the index finger of the dominant hand. The test involved two types of stimulus sequence, and the RT for each sequence was used as a measure of performance. One sequence (*i.e.,* blue, blue, green, green, red, yellow, red, blue, green, yellow, blue) was presented twice during the first and final trials (Fig. 2a) and used to evaluate improvement in general learning of the sequential information. The other sequence (*i.e.,* green, green, blue, green, yellow, yellow, blue, red, yellow, red, green) was repeated 30 times in succession during the trials (Fig. 2a), and good performance in the middle trials indicated improvement in processing speed resulting from repeated learning of the sequencial information. Each sequence included the same total numbers of vertical, horizontal, and diagonal finger movements. There was no time limit for completion of the test, and the sequence was visible throughout the duration of the trial. A black line was presented beneath the rectangle position to provide a visual cue indicating which button should be pressed, and the test did not proceed until the participant had pressed the correct button. Participants’ eye movements were not restricted. Each trial was followed by visual feedback lasting 3,000 ms, which showed the RT, number of errors the participant had made during the trial, and the number of trials completed. To avoid muscle fatigue, the inter-trial interval (ITI) lasted 5,000 ms (Fig. 2c). During the feedback and ITI, participants were instructed to place their dominant hand beside the response pad.

A modified version of a delayed matching-to-sample task developed by Pinal et al.^31^ was used to measure working memory. The test consisted of 100 trials, each beginning with an ITI of 800 ms, which reminded participants of the number of trials they had completed. After 500 ms, the encoding stimulus was presented for 1,000 ms (Fig. 2d), followed by a maintenance period of 5,000 ms and presentation of the recognition stimulus for 3,000 ms or until the participant provided a response. To restrict participants’ eye movements as much as possible, a fixation cross was presented at the centre of the monitor when no stimuli were present. The encoding and recognition stimuli consisted of squares of equal size (9.13° × 9.13° visual angle), and each square consisted of nine small coloured squares of equal size, as in each side of the standard Rubik’s Cube (Fig. 2d), produced via a custom-made MATLAB programme (Version R2015a, MathWorks Inc., Massachusetts, USA). The small squares were white, red, orange, yellow, green, or blue, consistent with the colours of the sides of the standard Rubik’s Cube. In most trials, the encoding and recognition stimuli were identical with the exception of one small square that was selected pseudo-randomly; however, the colour arrangements were identical in a few trials. The participants were asked to memorize the arrangement of the coloured square used as the encoding stimulus, hold it in their minds during the maintenance period, then retrieve it and identify the position of the target square within the arrangement of the nine small squares, which differed from the encoding stimulus in terms of colour. The participants were instructed to press one of nine keys, which corresponded to the position of the target, as quickly and accurately as possible, using a numeric keypad (model BSTK08, BUFFALO INC, Aichi, Japan). When the colour arrangements for the encoding and recognition stimuli were identical, participants were instructed to press 0. This test was divided into two blocks with different difficulty levels (Fig. 2e). The easy, or low memory load, block consisted of 25 trials in which squares consisted of three colours, and 25 trials in which squares consisted of four colours. The difficult, or high memory load, block consisted of 25 trials in which squares consisted of five colours, and 25 trials in which squares consisted of six colours. Each block included 10 trials in which the encoding and recognition stimuli were identical. The order of the blocks was randomized between participants but remained the same for each participant’s pre- and post-training tests, and stimuli were presented randomly within each block. The stimulus sets differed between the pre- and post-training tests.

A modified version of the task previously introduced by Peters et al.^33^ was used as the mental rotation test, to assess participants’ visuospatial ability. The stimulus set, which was developed by Ganis and Kievit,^34^ included eight images (7.65° × 7.78° visual angle): one standard object, three objects rotated 50°, 100°, and 150° about the horizontal axis, and four objects replaced by mirror images corresponding to these objects.^34^ In each trial, an image of a set of stimuli was presented at the top of the screen as the target stimulus, and four images (two standard images and two mirror images of the target object) were presented at the bottom of the screen as probe stimuli (Fig. 2f). In total, 45 stimulus sets were presented, and the order of presentation was randomized. Participants were instructed to identify both standard images of the target using mouse clicks and provide as many correct responses as possible within 3 min. The subsequent trial began immediately after participants had chosen two images in the trial.

### Data analysis

In the serial reaction time test, RTs, which were expressed as the total time taken to complete the sequence, were obtained for each sequence by summing all RTs obtained via the 11 button presses. First, analyses were performed using normalized data (normalized to performance in the first trials of the pre- and post-training tests) to rule out potential confounding group differences. In the middle trials, changes in RTs across 30 repetitions were compared between the three groups. In addition, we assumed that the RT for any sequence, including the first and final trials, would decrease if an extended effect was caused by the middle trials. To test this assumption, we calculated the difference in RTs between the first and final trials. RT differences between the pre- and post-training tests were then compared between the three groups. Thereafter, to confirm that the results involving normalized RTs reflected the transfer effects of the training accurately, raw RTs were also analysed. In the middle trials, differences in RTs between the post- and pre-training tests were used to compare changes in RTs prior and subsequent to training between the three groups. In the analysis of the first and final trials, RTs for the first trial were subtracted from those for the final trial for each group and test. Only one participant in the NCG group was excluded from the analyses, as his RTs were identified as outlier values via multivariate outlier analysis using Mahalanobis distance.

In the working memory test, the proportion of correct responses (PC) in all trials and the mean RT for correct responses were calculated in each block. Both measures were combined using the IES, which consists of the mean RT divided by the PC as follows:

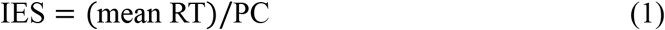

As RTs are measured in ms and divided into proportions, the IES is also measured in ms,^32^ and a higher IES reflects poorer performance. Furthermore, this index could be considered an ‘adjusted RT’ that prevents potential speed-accuracy trade-offs or criterion shifts. An IES was calculated for each participant and block. The differences in IESs between pre- and post-training tests were compared between the three groups.

In the mental rotation test, one point was awarded if both of the participant’s choices were correct. Therefore, the maximum score was 45. Pre- and post-training differences in scores were compared between the three groups.

## Statistical analyses

Statistical analyses were performed using JMP 12 (SAS Institute Inc., North Carolina, USA). Student’s *t* test was performed to examine differences in overall training time between the two training groups. In addition, two-way, repeated measures ANOVA (Group [CG, NCG] × Cube [familiar, unfamiliar]) was performed to compare the periods required to solve the Rubik’s Cube puzzle between the two training groups and between the training groups and the control group. Changes in motivation scores during the training session were compared between the three groups via a two-way ANOVA (Group [CG, NCG, control] × Week [1, 2, 3]). In the serial reaction time test, the reduction of normalized RTs in the middle trials was assessed separately for the pre- and post-training tests via a two-way, repeated measures ANOVA (Group [CG, NCG, control] × Trial [30 repetitions]). Thereafter, the normalized RT for the final trial was compared between the three groups via a one-way ANOVA (Group [CG, NCG, control]). A two-way, repeated measures ANOVA (Group [CG, NCG, control] × Trial [30 repetitions]) was performed to analyse raw RTs in the middle trials. Student’s *t* test was used to analyse raw RTs in the first and final trials for each group and test. Regarding the working memory test, the difference in pre- and post-training IESs was examined using a two-way ANOVA (Group [CG, NCG, control] × Load [low, high]). Differences in pre- and post-training scores in the mental rotation test were analysed via a one-way ANOVA (Group [CG, NCG, control]). Post-hoc Bonferroni-correction was performed for all analyses where necessary.

## Data availability

The data that support the findings of this study are available from the corresponding author upon reasonable request.

## Code availability

The custom computer codes that support the findings of this study are available from the corresponding author upon reasonable request.

## Acknowledgements

We would like to thank M. Hamakawa, N. Tsuzuki, and T. YuHsuan for their helpful suggestions and comments. The study was supported by the Qdai-jump Research Programme (27818, 2015-2016).

## Author contributions

M.S. designed the study while actively discussing it with the co-authors, K.T. and T.O.

M.S conducted the experiments, performed the data analysis, and wrote the paper. All of the authors discussed the results and revised the manuscript.

## Additional information

Supplementary information is available for this paper.

